# Study of Sex Differences in the Whole Brain White Matter Using Diffusion MRI Tractography and Suprathreshold Fiber Cluster Statistics

**DOI:** 10.1101/2025.09.27.679006

**Authors:** Fan Zhang, Jarrett Rushmore, Yijie Li, Suheyla Cetin-Karayumak, Yang Song, Weidong Cai, Carl-Fredrik Westin, James J. Levitt, Nikos Makris, Yogesh Rathi, Lauren J. O’Donnell

## Abstract

Sex-specific characteristics demonstrate a substantial influence on the brain white matter (WM), suggesting distinct structural connectivity patterns between females and males. Diffusion MRI (dMRI) tractography is an important tool in assessing WM connectivity and brain tissue microstructure across different populations. Whole brain tractography analysis for group statistical comparison is a challenging task due to the large number of WM connections. This work studies whole-brain WM connectivity differences between females and males using dMRI tractography. We study a large cohort of 707 subjects from the Human Connectome Project Young Adult dataset. By applying a well-established fiber clustering pipeline and a suprathreshold fiber cluster statistical method, we analyze tracts in the cerebral cortex and understudied pathways like those connecting to the cerebellum. We identify several tracts with significant sex differences in terms of their fractional anisotropy and/or mean diffusivity. These include deep tracts like the arcuate fasciculus, corticospinal tract, and corpus callosum, superficial tracts in the frontal lobe, and cerebellar tracts. Finally, correlation analysis reveals that these WM differences are linked to a range of neurobehavioral measures, with the strongest and most consistent associations observed for motor function, suggesting motor circuits as a potential key focus for future research.

## 1. Introduction

Sex-specific characteristics demonstrate a substantial influence on the human brain (Baron-Cohen et al., 2005; Cahill, 2006; Halpern, 2000). There has been an enduring interest in investigating neuroanatomical evidence of sexual dimorphism and its relationship to sex-specific behaviors (Giedd et al., 2012; Peper et al., 2011; Ruigrok et al., 2014). In particular, there is an evident prevalence of sex-specific differences in the brain white matter, suggesting distinct brain structural connectivity patterns between females and males (de Vries & Södersten, 2009; G. Gong et al., 2011). Diffusion magnetic resonance imaging (dMRI) (Basser et al., 1994) is the only existing non-invasive technique to map the brain white matter connections. dMRI measures the random molecular motion or diffusion of water molecules, allowing estimation of white matter fiber tracts in the brain via a process called tractography (Basser et al., 2000) and quantification of the microstructure of the white matter connections (Basser, 1995). Many studies have investigated white matter sexual dimorphism using dMRI tractography (Choi et al., 2010; Clayden et al., 2012; Eluvathingal et al., 2007; Hasan et al., 2009; Kitamura et al., 2011; Lebel et al., 2010; Lei et al., 2016; F. Liu et al., 2010; Y. Liu et al., 2011; Nordahl et al., 2015; Oh et al., 2007; Ritchie et al., 2018; Thiebaut de Schotten et al., 2011; G. Wang et al., 2014; J. Zhang et al., 2014); (Daianu et al., 2012; Dennis et al., 2013; G. Gong et al., 2009; Herlin et al., 2024; Ingalhalikar et al., 2014; Jahanshad et al., 2011; Tyan et al., 2017; Wheelock et al., 2021; Yan et al., 2011). Among these, most existing studies have applied the traditional tract of interest analysis to investigate differences in specific anatomical tracts without considering the entire white matter (Choi et al., 2010; Clayden et al., 2012; Eluvathingal et al., 2007; Hasan et al., 2009; Herlin et al., 2024; Kitamura et al., 2011; Lebel et al., 2010; Lei et al., 2016; F. Liu et al., 2010; Y. Liu et al., 2011; Nordahl et al., 2015; Oh et al., 2007; Ritchie et al., 2018; Thiebaut de Schotten et al., 2011; G. Wang et al., 2014; Wheelock et al., 2021; J. Zhang et al., 2014). On the other hand, there is an increased interest in whole brain tractography analysis to assess sex-specific brain structural connectivity patterns across the entire brain (Daianu et al., 2012; Dennis et al., 2013; G. Gong et al., 2009, 2011; Ingalhalikar et al., 2014; Jahanshad et al., 2011; Tyan et al., 2017; Yan et al., 2011). In the present work, we propose to study sex-specific differences in the whole brain white matter connections using dMRI tractography.

Whole brain tractography analysis requires a parcellation of the tractography data to enable simultaneous analysis across a large number of white matter connections in the entire brain (F. Zhang et al., 2022). Performing whole brain tractography on one individual subject can generate hundreds of thousands of streamlines, which are not immediately useful for quantitative statistical analysis. Tractography parcellation is thus needed to divide the massive number of tractography streamlines into anatomically meaningful fiber tracts (bundles or fascicles). Currently, whole brain tractography studies on sex differences have focused on the traditional connectome-based analysis, which parcellates tractography based on white matter connections between brain cortical regions, enabling the study of the brain using graph theory (Daianu et al., 2012; Dennis et al., 2013; G. Gong et al., 2009; Ingalhalikar et al., 2014; Jahanshad et al., 2011; Tyan et al., 2017; Yan et al., 2011). Nevertheless, there are several known challenges in the application of connectome-style analyses to dMRI (Sotiropoulos & Zalesky, 2019), including anatomical variability of white matter tract terminations and cortical anatomy (Bajada et al., 2017; Hau et al., 2016), spatial distortions (Jones & Cercignani, 2010), false positive tracking (Maier-Hein et al., 2017), and reduced reproducibility due to the non-continuous nature of the matrix representation (Moyer et al., 2017).

Another popular method to perform whole brain tractography analysis applies the fiber clustering strategy (O’Donnell et al., 2013), which parcellates tractography by grouping the streamlines according to their geometric trajectories. Unlike traditional connectome analysis, which is gray-matter-centric and studies graph properties of the connectivity matrix between cortical regions, fiber clustering analysis is white-matter-centric and enables the study of the microstructural characteristics of brain connections with a focus on white matter anatomy. Fiber clustering has been shown to have several advantages compared to the cortical-parcellation-based method, including highly consistent tract identification across populations (Cetin-Karayumak et al., 2024; Ge et al., 2012; Li et al., 2024; F. Zhang et al., 2017; F. Zhang, Wu, Norton, et al., 2018; Ziyan et al., 2009), highly reproducible tractography parcellation (Kai & Khan, 2019; F. Zhang et al., 2019), robustness to the presence of false positive streamlines (Jin et al., 2014; Sydnor et al., 2018), and improved power for predicting human traits (R. Liu et al., 2023). Fiber clustering has been widely used to investigate the brain’s structural connectivity in various applications in health and disease (Fan et al., 2019; Lo et al., 2025; O’Donnell et al., 2017; Stojanovski et al., 2019; Tunç et al., 2016; Wu et al., 2018; F. Zhang, Savadjiev, et al., 2018), providing a useful tool to investigate white matter sex differences.

In this work, we propose to investigate sex differences in the whole brain white matter using dMRI tractography fiber clustering. We leverage a well-established data-driven fiber clustering white matter parcellation pipeline (Li et al., 2024; O’Donnell et al., 2012; O’Donnell & Westin, 2007; F. Zhang, Wu, Norton, et al., 2018), which has been successfully applied in multiple neuroscientific studies including brain disorder analysis (Irimia et al., 2020; Levitt et al., 2023; Wu et al., 2018; F. Zhang, Savadjiev, et al., 2018; F. Zhang, Wu, Ning, et al., 2018), neurosurgical planning research (S. Gong et al., 2018; O’Donnell et al., 2017; F. Zhang, Noh, et al., 2020), and brain atlasing (F. Zhang, Wu, Norton, et al., 2018; F. Zhang, Xie, et al., 2020). One of the advantages of the fiber clustering method is that, together with our white matter atlas (F. Zhang, Wu, Norton, et al., 2018), it enables a fine-grained whole brain tractography parcellation into a total of 1516 clusters. These clusters include not only fiber connections in the deep white matter but also superficial and cerebellar white matter connections that have been relatively less studied in previous work about sex differences. In addition, we utilize an advanced suprathreshold fiber cluster (STFC) statistical analysis method (F. Zhang, Wu, Ning, et al., 2018), specifically designed for the fiber clustering pipeline, for testing group differences in whole brain tractography. STFC enables simultaneous statistical group comparison across a large number of white matter connections. It leverages permutation testing and whole brain white matter geometry to enhance the statistical analysis of tractography while correcting for multiple comparisons. The STFC method has been successfully applied to study group differences in attention-deficit/hyperactivity disorder (ADHD) and healthy controls (F. Zhang, Wu, Ning, et al., 2018), showing a highly sensitive ability to detect white matter group differences.

We perform the study on a large healthy adult cohort (n = 707) from the Human Connectome Project Young Adult (HCP-YA) (Van Essen et al., 2013). The HCP-YA acquired high-quality *in vivo* macroscopic-level imaging data in an effort to elucidate the neural pathways and networks that underlie brain function and behavior (Barch et al., 2013). Non-imaging demographics (age, sex, etc.) and behavioral (fluid intelligence, language performance, etc.) measures are also available. Many research studies have leveraged the HCP-YA data and revealed important findings related to the human brain (Liégeois et al., 2019; Smith et al., 2015), including sex differences in brain gray matter and white matter volumes using T1-weighted MRI (Szalkai & Grolmusz, 2018) and in brain functional connectivity using functional MRI (Sen & Parhi, 2019; Weis et al., 2019; C. Zhang et al., 2018). A recent study utilized dMRI data to examine sex differences in major deep white matter fiber tracts using the HCP-YA cohort (Herlin et al., 2024). In our study, we aim to build upon this work by analyzing white matter connections across the entire brain, including the deep and superficial white matter, and further exploring their relationship to behavioral measures.

In the rest of this paper, we first describe the datasets under study, including dMRI data from a large cohort of 707 healthy adult subjects from the HCP-YA (Van Essen et al., 2013). Then, we introduce the related computational processes, including multi-tensor tractography and fiber clustering white matter parcellation, followed by the extraction of diffusion features of interest (fractional anisotropy (FA) and mean diffusivity (MD)). Next, we demonstrate the process of STFC statistical comparison to identify the white matter fiber clusters that are significantly different between the female and male groups. Following that, a correlation analysis is performed to investigate the relationship between the diffusion measures of the significant fiber clusters and the behavioral measures for each of the female and male groups. Finally, we give the experimental results and related discussion.

## 2. Methods

### 2.1. Datasets

We included a large cohort of 707 subjects (340 females and 367 males) in our study, with the following subject selection procedures from the entire HCP-YA population. (1) We excluded the 100 subjects that were used to generate the fiber clustering white matter atlas (F. Zhang, Wu, Ning, et al., 2018) (see Section 2.3 for a brief introduction of the atlas). This exclusion was to avoid any potential bias in the tractography parcellation across the subjects under study. (2) We excluded one of a twin pair if both twins participated in the HCP-YA study. (3) We selected the maximal subset of subjects that had a statistically similar age distribution between the male and female groups. In total, we selected 707 subjects, including 340 females and 367 males (mean age: 28.2±3.1y versus 28.0±3.6y, p=0.43).

The HCP-YA dMRI data was acquired with a high-quality image acquisition protocol using a customized Connectome Siemens scanner (Van Essen et al., 2012) and was processed following the well-designed HCP-YA minimum processing pipeline (Glasser et al., 2013). The acquisition parameters for the dMRI data were TE = 89.5 ms, TR = 5520 ms, phase partial Fourier = 6/8, voxel size = 1.25 x 1.25 x 1.25 mm^3^, 18 baseline images with a low diffusion weighting b = 5 s/mm^2^, and 270 diffusion-weighted images evenly distributed at three shells of b = 1000/2000/3000 s/mm^2^. More detailed information about the HCP-YA data acquisition and preprocessing can be found in (Glasser et al., 2013).

### 2.2. Multi-tensor whole brain tractography

Whole brain tractography was computed for each subject under study using the two-tensor Unscented Kalman Filter (UKF) method (Malcolm et al., 2010; Reddy & Rathi, 2016), as implemented in the *ukftractography* package (https://github.com/pnlbwh/ukftractography). The UKF method fits a mixture model of two tensors to the dMRI data while tracking, employing prior information from the previous step to help stabilize model fitting. The mixture model assumes there are two distinct tract directions within voxels, in which the first one represents the principal direction and the other one is from the second tract. This multi-tensor approach enables streamline-specific microstructure estimation. We chose the UKF method because it has been shown to be highly consistent in tracking streamlines in dMRI data from independently acquired populations across ages, health conditions, and image acquisitions (F. Zhang, Wu, Ning, et al., 2018), and it is more sensitive than standard single-tensor tractography, in particular in the presence of crossing fibers and peritumoral edema (Baumgartner et al., 2012; Chen et al., 2015, 2016; Liao et al., 2017). In addition, we chose the UKF method because it was used to generate tractography data in the fiber clustering atlas (F. Zhang, Wu, Ning, et al., 2018) (see Section 2.3 for a brief introduction of the atlas). Using the same tractography method can ensure highly consistent tract parcellation results across subjects (F. Zhang et al., 2017; F. Zhang, Wu, Ning, et al., 2018). We adopted the same tractography parameters as used in (F. Zhang, Wu, Ning, et al., 2018), producing about 1 million streamlines per subject. Visual and quantitative quality control of the tractography was performed using a quality control tool in the *whitematteranalysis* (WMA) software (https://github.com/SlicerDMRI/whitematteranalysis).

### 2.3. Tractography fiber clustering

Fiber clustering was performed to parcellate the whole brain tractography data for each individual subject using a well-established fiber clustering pipeline (O’Donnell et al., 2012; O’Donnell & Westin, 2007) and a fiber clustering atlas provided by the O’Donnell Research Group (ORG) (F. Zhang, Wu, Norton, et al., 2018). The ORG atlas contains an 800-cluster parcellation of the entire white matter and an anatomical fiber tract parcellation (http://dmri.slicer.org/atlases)^1^. The atlas was generated by creating dense tractography maps (using the same UKF tractography method as in the current study) of 100 individual HCP-YA subjects and then applying a fiber clustering method to group the tracts across subjects according to their similarity in shape and location. The resulting clusters were annotated using expert neuroanatomical knowledge. In the present study, we used the 800-cluster parcellation for whole brain white matter connectivity analysis, while the anatomical fiber tract parcellation was used to evaluate to which anatomical structure the identified significantly different white matter connections belonged.

The fiber clustering method was applied to perform white matter parcellation of one subject as follows. A tractography-based registration was performed to align the subject’s tractography data into the atlas space. A fiber spectral embedding was conducted to compute the similarity of streamlines between the subject and the atlas, followed by the assignment of each streamline of the subject to the corresponding atlas cluster. This process produced a whole brain white matter parcellation into 800 clusters. These clusters included 84 commissural clusters as well as 716 bilateral hemispheric clusters (that included streamlines in both hemispheres). We separated the hemispheric clusters by hemisphere, thus resulting in (716 x 2 + 84) = 1516 clusters per subject. All fiber clustering processing was performed using the *whitematteranalysis* software (https://github.com/SlicerDMRI/whitematteranalysis), and all parameters were set to their default values. (Details are described in (O’Donnell et al., 2012, 2017; O’Donnell & Westin, 2007; F. Zhang, Wu, Norton, et al., 2018).)

### 2.4. Quantitative diffusion measure computation

Diffusion measures were extracted from each fiber parcel to quantify the microstructural properties of the brain white matter connections. In this study, we focused on the FA and MD measures. FA and MD are quantitative microstructure measures derived from modeling water diffusion using dMRI mathematics (Basser, 1995). They are sensitive to the differences in the underlying cellular microstructure in brain tissues. Studies in the literature have observed differences in FA and/or MD in multiple white matter connections between females and males (Choi et al., 2010; Eluvathingal et al., 2007; Kitamura et al., 2011; Lebel et al., 2010; F. Liu et al., 2010; Y. Liu et al., 2011; Oh et al., 2007; Thiebaut de Schotten et al., 2011). Specifically, in the present study, we computed the mean FA and the mean MD of each fiber parcel.

### 2.5. STFC statistical comparison for group differences

For each diffusion measure, statistical group comparison was performed using the STFC method (F. Zhang, Wu, Ning, et al., 2018). The STFC method leverages permutation testing and whole brain white matter geometry to enhance the statistical analysis of tractography. The basic idea of the STFC method is similar to the well-known voxel-cluster-thresholding approach (Nichols & Holmes, 2002; Smith & Nichols, 2009), but with the improvement of leveraging a fiber parcel’s neighborhood (nearby parcels with similar white matter anatomy) to support each parcel’s statistical significance when correcting for multiple comparisons.

The STFC method employs several steps, as described in detail in the original publication (F. Zhang, Wu, Ning, et al., 2018). Here, we give a brief overview of the steps of the STFC method as employed in this study. Specifically, for each diffusion measure (FA or MD), the STFC method first analyzed each fiber cluster using a general linear model (GLM) to compare between the female and male groups, with age as a covariate to control for potential age differences. Next, fiber clusters with uncorrected p<0.05 from the GLM analysis were identified. Then, these identified fiber clusters were combined with their neighbors to form STFCs, which contained multiple neighboring clusters, where neighborhoods were defined based on streamline geometric similarity. Finally, a nonparametric permutation test was used to determine a corrected p-value for each STFC based on its STFC size, i.e., the number of fiber clusters within the STFC, under the null hypothesis of no group difference in any cluster. The STFCs with a corrected p<0.05 were considered to be significantly different between the female and male groups.

### 2.6. Canonical correlation analysis of behavioral measures

We then investigated the relationship between the diffusion measures of the identified STFCs and the behavioral measures for each of the female and male groups. We adopted the sophisticated multivariate analysis method proposed in (Smith et al., 2015), which was designed to study the relationship between functional MRI connectomes and behavioral data in HCP. This method uses canonical correlation analysis (CCA) (Hotelling, 1936) to identify pairs of canonical variates along which sets of behavioral measures and patterns of imaging measures co-vary in a similar way across subjects. Each pair of variates is referred to as a mode of co-variation, associated with a coefficient *r* that indicates the strength of the canonical correlation.

We investigated the behavioral measures belonging to the cognition, emotion, motor, personality, and sensory categories, respectively, provided in the HCP-YA database (as documented in HCP-YA S1200 Data Dictionary). For each diffusion measure (FA or MD), we performed two CCA analyses to assess relationships between the behavioral measures and the diffusion measures of the significant STFCs in the female and male groups, respectively. Thus, for each behavioral category, there were a total of 4 CCA analyses across the two diffusion measures and two groups. For each CCA analysis, the major mode (i.e., the one with the highest coefficient *r*) was reported. A permutation test was performed to determine the statistical significance of the major mode by testing for a null distribution of *r* via 100,000 permutations of the behavioral measures relative to the diffusion measures (Smith et al., 2015). A p-value lower than 0.05 (FDR corrected for multiple comparisons across the four modes) was considered statistically significant.

## 3. Results

### 3.1. Sex differences in white matter FA values

The STFC identified nine white matter regions that exhibited significant sex differences in fractional anisotropy Table 1; Figure 1). These differences were observed in several white matter pathways that showed a significantly larger FA in females: coronal radiata bilaterally, middle and inferior cerebellar peduncles, and right thalamo-temporal projections. Males exhibited a larger FA in the superficial frontal white matter bilaterally and in the left thalamo-frontal projection.

**Figure 1.**
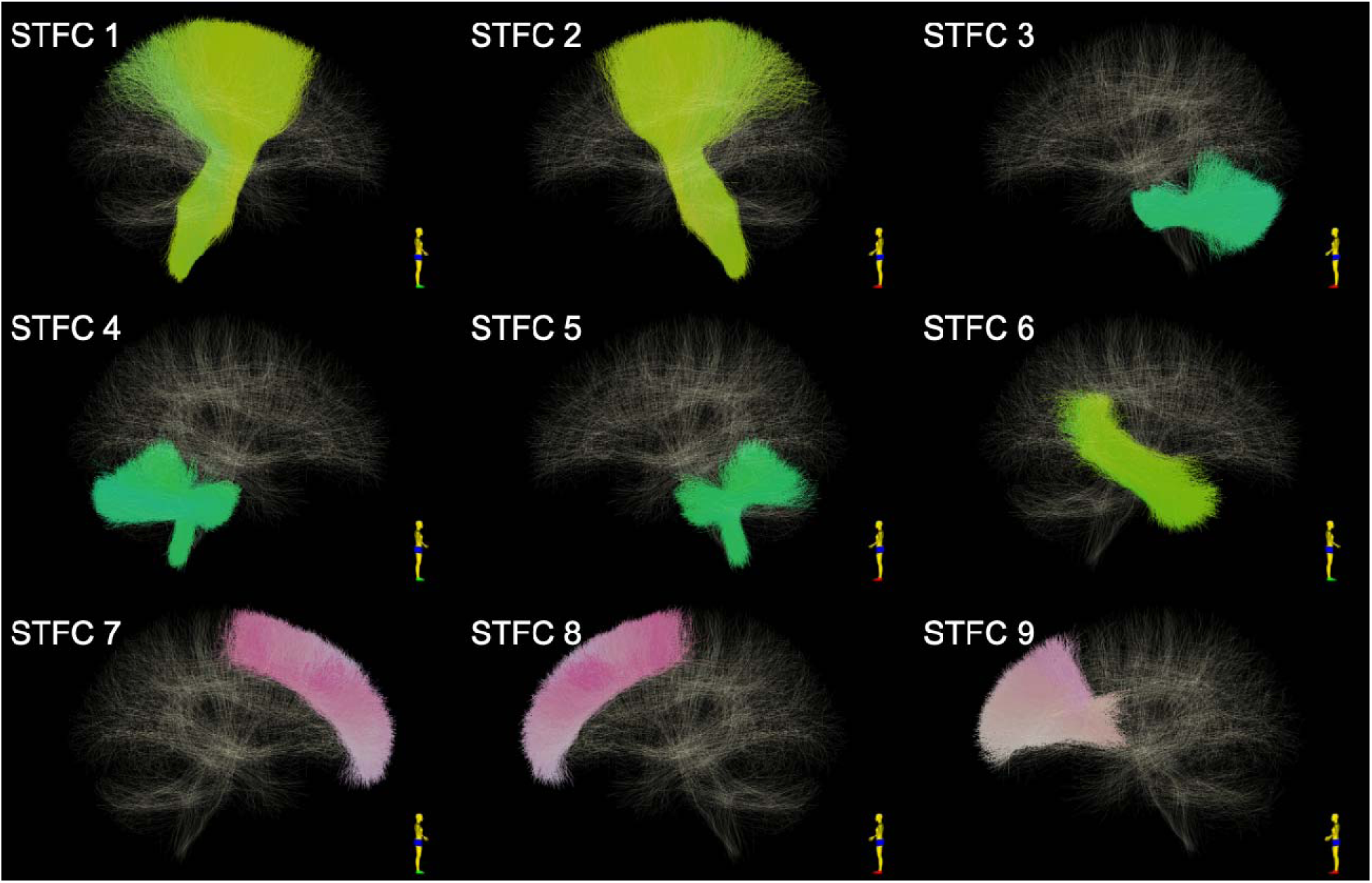
Visualization of the STFCs that are identified to be significantly different between females and males using FA.

**Table 1.** Significant differences in white matter regions between females and males using FA.

| STFC index | STFC size | Anatomical structure | p-value | Mean FA of clusters in STFC |  |
| --- | --- | --- | --- | --- | --- |
| | | | | Female group (Mean±SD) | Male group (Mean±SD) |
| STFC 1 | 11 | Corona radiata (right) | 0.0105 | **0.6431 ± 0.0208** | 0.6350 ± 0.0218 |
| STFC 2 | 13 | Corona radiata (left) | 0.0036 | **0.6333 ± 0.0204** | 0.6246 ± 0.0215 |
| STFC 3 | 8 | Middle cerebellar peduncle | 0.0422 | **0.5213 ± 0.0235** | 0.5100 ± 0.0272 |
| STFC 4 | 13 | Middle/inferior cerebellar peduncle (right) | 0.0036 | **0.4021 ± 0.0234** | 0.3856 ± 0.0303 |
| STFC 5 | 8 | Middle/inferior cerebellar peduncle (left) | 0.0422 | **0.4004 ± 0.0256** | 0.3773 ± 0.0308 |
| STFC 6 | 8 | Thalamo-temporal (right) | 0.0422 | **0.3695 ± 0.0222** | 0.3614 ± 0.0243 |
| STFC 7 | 15 | Superficial frontal (right) | 0.0012 | 0.3894 ± 0.0269 | **0.4011 ± 0.0259** |
| STFC 8 | 15 | Superficial frontal (left) | 0.0012 | 0.3849 ± 0.0261 | **0.3970 ± 0.0266** |
| STFC 9 | 12 | Thalamo-frontal (left) | 0.0062 | 0.3821 ± 0.0245 | **0.3976 ± 0.0268** |

### 3.2. Sex differences in white matter MD values

The STFC approach identified seven white matter regions that exhibited significant sex differences in mean diffusivity (Table 2; Figure 2). Females exhibited lower MD in the coronal radiata bilaterally, the left arcuate fascicle, and the corpus callosum (subdivisions 4 to 7). By contrast, males had lower MD in the left intracerebellar parallel tract, the right middle/inferior cerebellar peduncle, and the right thalamo-temporal projection.

**Figure 2.**
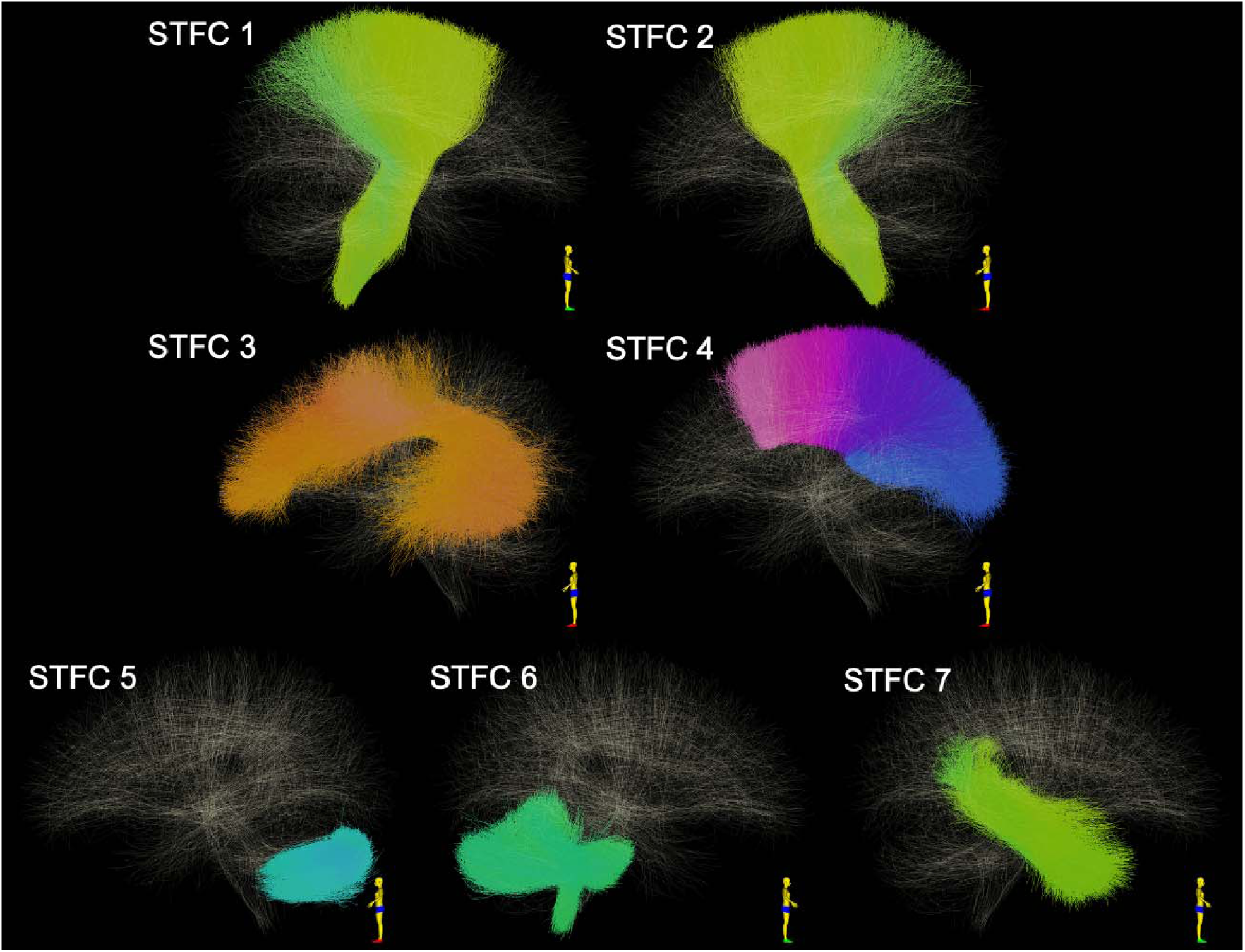
Visualization of the STFCs that are identified as significantly different between females and males using MD.

**Table 2.** Significant differences in white matter regions between females and males using MD.

|  |  |  |  |  |
| --- | --- | --- | --- | --- |
| STFC | STFC | Anatomical structure | p-value | Mean MD of clusters in STFC ($10^{-3}$) |

| index | size |  |  | Female group<br>(Mean±SD) | Male group<br>(Mean±SD) |
| --- | --- | --- | --- | --- | --- |
| STFC 1 | 12 | Corona radiata (left) | 0.0263 | **0.1303 ± 0.0027** | 0.1288 ± 0.0027 |
| STFC 2 | 12 | Corona radiata (right) | 0.0263 | **0.1297 ± 0.0026** | 0.1280 ± 0.0027 |
| STFC 3 | 12 | Arcuate fasciculus (left) | 0.0263 | **0.1276 ± 0.0034** | 0.1258 ± 0.0035 |
| STFC 4 | 18 | Corpus callosum (4-7) | 0.0002 | **0.1179 ± 0.0030** | 0.1159 ± 0.0029 |
| STFC 5 | 12 | Intracerebellar parallel tract (left) | 0.0263 | 0.1371 ± 0.0037 | **0.1389 ± 0.0044** |
| STFC 6 | 12 | Middle/inferior cerebellar<br>peduncle (right) | 0.0263 | 0.1496 ± 0.0035 | **0.1513 ± 0.0037** |
| STFC 7 | 20 | Thalamo-temporal (right) | 0.0001 | 0.1329 ± 0.0036 | **0.1358 ± 0.0042** |

### 3.3. Canonical correlation with behavioral metrics

Canonical correlation analyses were performed to investigate the relationship between the white matter microstructure and behavior in females and males. Analysis was performed separately for FA and MD, and five behavioral domains (cognition, emotion, motor, personality, sensory) from the HCP-YA dataset were evaluated (Table 3). In the female group, FA values in the significant STFC were correlated with motor performance and sensory processing, while in males, FA was correlated only with motor performance. For MD, significant correlations were identified in the female group with cognitive performance and motor behavior, whereas in males, MD was again significantly correlated with motor performance.

**Table 3:** CCA analysis between STFC diffusion measures and subject behavioral measures. Correlation coefficient and p-value (permutation testing, with FDR correction) are reported.

|  |  | Cognition | Emotion | Motor | Personality | Sensory |
| --- | --- | --- | --- | --- | --- | --- |
| FA | Female | 0.421 (p=0.082) | 0.400 (p=0.361) | 0.321 (**p=0.015**) | 0.268 (p=0.259) | 0.412 (**p=0.002**) |
|  | Male | 0.395 (p=0.157) | 0.378 (p=0.433) | 0.301 (**p=0.016**) | 0.256 (p=0.259) | 0.301 (p=0.259) |
| MD | Female | 0.429 (**p=0.021**) | 0.416 (p=0.101) | 0.289 (**p=0.025**) | 0.187 (p=0.867) | 0.290 (p=0.237) |
|  | Male | 0.363 (p=0.237) | 0.360 (p=0.469) | 0.304 (**p=0.006**) | 0.241 (p=0.237) | 0.300 (p=0.136) |

## 4. Discussion

In this study, we investigated sex differences in the whole brain white matter using dMRI tractography fiber clustering. We performed whole brain fiber tracking using a multi-fiber UKF tractography method (Malcolm et al., 2010; Reddy & Rathi, 2016). We leveraged a well-established data-driven fiber clustering pipeline (O’Donnell et al., 2012; O’Donnell & Westin, 2007) and an anatomically curated white matter atlas (F. Zhang, Wu, Norton, et al., 2018) for tractography parcellation. We extracted two diffusion measures of interest (FA and MD) for the quantification of the tissue microstructure of each fiber cluster. We adopted the advanced STFC method for statistical between-group comparison (F. Zhang, Wu, Ning, et al., 2018). A large population, including a total of 707 healthy adult subjects (340 females and 367 males) from the HCP-YA database, was studied. Overall, we found sex differences in the whole brain white matter connectivity of the human brain.

We have several specific observations regarding the comparison results between the female and male groups under study. First, we found multiple significantly different white matter connections from the entire brain (a total of 16 STFCs). These white matter connections included several deep white matter tracts (e.g., arcuate fasciculus, corticospinal tract, and corpus callosum) that have been previously shown to have sex differences (see below for related work and detailed discussion). We also identified fiber tracts with significant sex differences from the superficial and the cerebellar white matter regions that have been relatively less investigated before using dMRI tractography. Second, there was no consensus on the direction of differences in FA and MD across the multiple identified STFCs. Nine STFCs were identified using FA, of which 4 had significantly higher FA in the female group and 3 had significantly higher FA in the male group. Six STFCs were identified using MD, of which 4 had significantly higher MD in the female group and 2 had significantly higher MD in the male group. Third, with the CCA analysis between the tract microstructure measures and the behavioral data, we found that the identified white matter connections were related to multiple brain functions, including motor, sensory, and cognitive functions. Significant correlations with motor function were present in both groups, while identified connections showed significant correlations with cognition and sensory processing in the female group and sensory processing in the male group.

Numerous studies have applied dMRI to study sex-specific differences in the brain’s white matter. The first category of studies has focused on voxel-based analysis. Traditional voxel-based dMRI studies have been conducted using region-of-interest (ROI) methods to identify white matter differences in a local region (Szeszko et al., 2003). Other voxel-based studies have performed whole brain white matter analysis using approaches such as voxel-based morphometry (VBM) (Dunst et al., 2014) and tract-based spatial statistics (TBSS) (Bava et al., 2011; Y. Wang et al., 2012). These voxel-based studies have revealed substantial sex-specific neuroanatomical characteristics in the white matter. However, using voxel-based methods, accurate localization of between-population differences to specific white matter tracts can be difficult to assess (Bach et al., 2014; Davatzikos, 2004), especially since more than one tract may cross within a single voxel (Raffelt et al., 2015; F. Zhang, Wu, Ning, et al., 2018).

Another category of dMRI studies has applied tractography-based analysis to investigate white matter sex differences. Tractography allows estimation of the trajectories of fiber tracts (Pujol et al., 2015; Yamada et al., 2009) and thus measurement of microstructural white matter properties of fiber pathways for a more detailed investigation of specific subpopulations of fibers (Ciccarelli et al., 2008; Hagmann et al., 2007). Hypothesis-driven tractography studies using tract-of-interest approaches have identified sex differences in many individual white matter fiber tracts (Choi et al., 2010; Clayden et al., 2012; Eluvathingal et al., 2007; Hasan et al., 2009; Herlin et al., 2024; Kitamura et al., 2011; Lebel et al., 2010; Lei et al., 2016; F. Liu et al., 2010; Y. Liu et al., 2011; Nordahl et al., 2015; Oh et al., 2007; Ritchie et al., 2018; Thiebaut de Schotten et al., 2011; G. Wang et al., 2014; J. Zhang et al., 2014). Despite differences in methodology, our study’s findings are in agreement with several recent tract-based findings in the HCP-YA dataset, including sex differences of microstructural parameters in the corticospinal tract (part of the corona radiata region we identified), cerebellar connections, and thalamo-temporal radiations (Herlin et al., 2024). Another category of tractography-based studies, to which our work belongs, has performed whole brain white matter analysis (Daianu et al., 2012; Dennis et al., 2013; G. Gong et al., 2009, 2011; Ingalhalikar et al., 2014; Jahanshad et al., 2011; Tyan et al., 2017; Yan et al., 2011). The benefits of whole brain tractography analysis include the assessment of sex-specific brain structural connectivity patterns across the entire brain, instead of specific tracts of interest. Recent neuroimaging studies, including not only dMRI but also other imaging modalities such as structural T1-and T2-weighted MRI, functional MRI, and positron emission tomography (PET), have shown sex-related differences likely exist in the whole brain (see a review (G. Gong et al., 2011) and several more recent studies (Hu et al., 2013; Tomasi & Volkow, 2012; Xin et al., 2019)). Another benefit of whole brain tractography analysis is enabling the data-driven identification of fiber tracts that have group differences (F. Zhang, Savadjiev, et al., 2018).

In the related literature of whole brain tractography sex difference studies, the methodology of cortical-parcellation-based connectome analysis has been widely used. In connectome analysis for studies of sex differences, a matrix of connection strengths between cortical regions is created, enabling the study of the brain using graph theory (Daianu et al., 2012; Dennis et al., 2013; G. Gong et al., 2009; Ingalhalikar et al., 2014; Jahanshad et al., 2011; Tyan et al., 2017; Yan et al., 2011). For instance, Ingalhalikar et al. constructed a connectome matrix based on a cortical/subcortical parcellation including 95 regions of interest (ROIs) and the number of tractography streamlines between each pair of the ROIs (thus leading to a 95 x 95 matrix). Multiple connectome measures have been studied by comparing graph theoretical properties across sexes (e.g., modularity, a measure of how well the network can be delineated into groups (see (Ingalhalikar et al., 2014) for details). Other widely studied connectome measures include small-worldness, weighted network cost, and clustering coefficient (G. Gong et al., 2009; Tyan et al., 2017). While connectome-style analyses have been widely used to study sex-specific brain connectivity differences, there are several technical challenges, including anatomical variability of fiber tract terminations and cortical anatomy (Bajada et al., 2017; Hau et al., 2016), spatial distortions (Jones & Cercignani, 2010), and reduced reproducibility due to the non-continuous nature of the matrix representation (Moyer et al., 2017). In our work, we applied another whole brain white matter connectivity analysis, i.e., dMRI tractography fiber clustering. In contrast, fiber cluster analysis is white-matter-centric, leveraging information about the full course of a fiber tract, which is the anatomical basis for fiber tract definition (O’Donnell et al., 2013; Wassermann et al., 2016). Previous studies have indicated that white-matter-centric approaches benefit from low variability across subjects in terms of the white matter parcellations that can be defined (Chekir et al., 2017; Guevara et al., 2017; O’Donnell et al., 2017; F. Zhang et al., 2017), and highly consistent tract identification across populations (Ge et al., 2012; Sydnor et al., 2018; F. Zhang et al., 2017, 2019; F. Zhang, Wu, Norton, et al., 2018; Ziyan et al., 2009), and highly reliable and reproducible performance for white matter parcellation (F. Zhang et al., 2019).

Whole brain analysis is intended to identify which connections are more different, which requires enhanced statistical power. In our study, we used a data-driven whole brain white matter atlas that finely divides all input tractography into many regions (including a total of 800 fiber clusters) and hence allows the identification of potential local white matter group differences from the whole brain. In addition, we applied our suprathreshold fiber cluster (STFC) method that leverages the whole brain fiber geometry to enhance statistical group difference analyses (F. Zhang, Wu, Ning, et al., 2018). The method performs statistical analyses of tractography for group comparison while correcting for multiple comparisons to allow simultaneous analysis across the entire white matter.

The identified white matter tracts that had significant sex differences in our study were obtained using the multi-fiber UKF tractography. The UKF has been shown to be more sensitive in tracking through the regions in the presence of crossing fibers (S. Gong et al., 2018; He et al., 2023; Liao et al., 2017; Xie et al., 2020; F. Zhang, Wu, Norton, et al., 2018). This could reduce the well-known false negative tractography problem and enable the investigation of more white matter connections (e.g., the corticospinal tract and arcuate fasciculus) compared to traditional single-tensor DTI tractography (Duffau, 2014; Jeurissen et al., 2013; Nimsky, 2014). A few sex difference studies have applied multi-tensor/fiber tractography for tract-of-interest analysis (Benavidez et al., 2024; Clayden et al., 2012; Kang et al., 2025; Y. Liu et al., 2011; Ritchie et al., 2018).

Next, we discuss detailed observations regarding the results for each significantly different white matter connection. Multiple identified deep white matter fiber tracts correspond to white matter structures that have been previously implicated to have sex differences, while we have also identified several cerebellar tracts and superficial tracts that have been rarely reported in the existing studies.

Bilateral tracts from the corona radiata to the brain stem, including the CSTs, were significantly different between the female and male groups under study in both FA and MD. Multiple studies have reported sex-specific differences in these white matter regions (Bava et al., 2011; Hervé et al., 2009; Seunarine et al., 2016; Takao et al., 2014). The corona radiata and CST are known to be related to human sensory and motor functions, which potentially explain the sensory and motor correlations in our CCA analysis. It is interesting to note that both FA and MD were significantly higher in the female group, suggesting greater diffusion anisotropy and diffusivity compared to the male group. Bava et al have reported a similar finding in bilateral CSTs using a voxel-based TBSS analysis (Bava et al., 2011).

The posterior portion of the corpus callosum tract was found to have significant sex differences in MD but not in FA in our study. The corpus callosum is one of the most widely studied white matter structures in the literature of dMRI-based sex differences. Early studies have identified the differences in the size of the corpus callosum between sexes, in particular, that of the corpus callosum splenium (the posterior portion) (Bishop & Wahlsten, 1997; DeLacoste-Utamsing & Holloway, 1982). Multiple studies have also identified tissue microstructural differences in the corpus callosum using voxel-based dMRI analysis, mostly with the findings of lower diffusion anisotropy and/or higher mean diffusivity in females (Menzler et al., 2011; Shin et al., 2005; Westerhausen et al., 2003). Our results are in line with these studies by showing a higher MD in the female group; however, we did not find any differences in the corpus callosum using FA. Our identified corpus callosum tract was specifically from the CC4 to CC7 (Makris et al., 1999), connecting to the motor cortex (CC4), the somatosensory cortex (CC5), the superior parietal lobe (CC6), and the lateral occipital cortex (CC7).

We identified the left arcuate fasciculus to be significantly different, where the female group had a significantly higher MD. The left arcuate fasciculus is known to be related to human language function (Zekelman et al., 2022). Our results in the left arcuate fasciculus, but not in the contralateral side, are possibly related to the sex difference of left-hemispheric language lateralization that has been reported in many studies (Allendorfer et al., 2016; Catani et al., 2007). Several studies have reported sex differences in the left arcuate fasciculus, mostly using FA (Jung et al., 2019; Madhavan et al., 2014) (significantly lower FA in females). Nevertheless, in our study, no FA differences were identified in the arcuate fasciculus.

We found a significantly higher FA of the left striato-frontal tract in the male group. The striato-frontal structural connections are known to play important roles in behaviors such as reward-related processes and impulse control (Morris et al., 2016). Previous research has hinted at potential sex differences in the striato-frontal connections. For example, multiple voxel-wise studies have shown that males had stronger white matter connections in white matter regions and tracts that underlie striatal-frontal connections (Lei et al., 2016), which is in line with our finding that the male group has a higher FA. More recently, we and another group have also identified sex differences with a tractography technique by measuring topographical organization (F. Zhang, Vangel, et al., 2020) and tract volume (Lei et al., 2016) in striato-frontal white matter connections. While bilateral differences were identified in these two studies as well as in the aforementioned voxel-wise study (Lei et al., 2016), in the present study, we only identified significant differences in the left hemisphere.

We also found that multiple cerebellar white matter tracts were significantly different in both FA and MD. The cerebellum is important and related to multiple brain functions (Mormina et al., 2017; Travis et al., 2015). Compared to the deep white matter, sex differences of the cerebellar white matter are relatively less studied, in particular using dMRI. A few studies have identified microstructure differences between sexes in the cerebellar peduncle (Bede et al., 2014; Hosoki et al., 2023; Kanaan et al., 2012, 2014; Takao et al., 2014) using voxel-based dMRI analysis. One dMRI tractography study has identified sex differences in the structural connectome between the cerebellum and the cerebral cortex (Ingalhalikar et al., 2014). Our results provided additional evidence for sexual dimorphism in the cerebellar white matter. We showed that there were significant sex differences in the middle cerebellar peduncle tract, particularly in the right hemisphere where both FA and MD were significantly different. Another interesting finding is that we found intracerebellar parallel tract connections were significantly different in MD, which has not yet been reported. Intracerebellar parallel tract connections are important and shown to be linked with sensorimotor functions (Jörntell, 2017).

Bilateral superficial white matter tracts in the superior frontal lobes were found to be significantly different between the female and male groups. Unlike the deep white matter structures that have been extensively studied in different clinical and scientific applications, the superficial white matter is relatively less studied (Phillips et al., 2013; Shiino et al., 2017). In our study, we find that only FA showed sex differences in the superficial white matter, where the female group had lower FA.

An important aspect of the current study is that it was performed in sufficient numbers to assess not only the fiber tracts that are significantly different between males and females, but also those fiber bundles that did not exhibit sex differences. Only a few short and long-range cortico-cortical association bundles differed on the basis of sex, suggesting limited differences between females and males. The superficial white matter of the frontal lobe had a higher FA in males, whereas the left arcuate fasciculus and the corpus callosum exhibited higher MD in females. FA and MD in the short association cortico-cortical white matter systems, such as the superficial white matter in the temporal, occipital, and parietal areas, did not vary according to sex., Moreover, most long-range cortico-cortical fiber bundles did not differ with sex (i.e., cingulum bundle, external capsule, extreme capsule, ILF, iFOF, MDLF, SLF, UF), suggesting that sex differences in cortical processing are limited.

Examination of cerebellar tracts revealed considerable sex differences, with the middle cerebellar peduncles being consistently affected. When taken together with the findings from the cerebral cortex, the most consistent findings may be related to motor function. Motor function and planning are largely localized to the frontal lobe, and bilateral differences in the frontal lobe superficial white matter, and the corticospinal and corticopontine tracts pass through the coronal radiata. In addition, the middle cerebellar peduncle, which carries signals from the corticopontine tract after relay in the ipsilateral pontine nucleus, relays motor signals to the cerebellum. These findings are consistent with the correlation analyses that show the significantly different tracts are most consistently related to neurobehavioral indices of motor function.

Our findings are broadly consistent with investigations using dMRI tractography in adolescents, which have shown males to have a larger within hemispheric connectivity, but lower cerebellar connectivity (Ingalhalikar et al., 2014).

## 5. Conclusion

We studied whole-brain white matter sex differences using a data-driven fiber clustering pipeline, a white matter parcellation atlas, and an advanced STFC statistical analysis method. We compared FA and MD between the two groups in a large cohort of 707 subjects. We identified multiple fiber tracts that had significant differences, which provides evidence pointing to sex-specific brain white matter connectivity.

## Data and Code Availability

The data under study are from the public Human Connectome Project (HCP) dataset (www.humanconnectome.org). The UKF code to perform fiber tracking is publicly available (https://github.com/pnlbwh/ukftractography). The WMA code to perform fiber clustering is publicly available (https://github.com/SlicerDMRI/whitematteranalysis). The STFC code to perform the statistical analysis is publicly available (https://github.com/zhangfanmark/STFC). Tract visualization is performed using SlicerDMRI (https://dmri.slicer.org).

## Author Contributions

Fan Zhang: Conceptualization, Data curation, Investigation, Project administration, Software, Visualization, Writing – original draft, Writing – review & editing

Jarrett Rushmore: Investigation, Result Interpretation, Writing – original draft, Writing – review & editing

Yijie Li: Investigation, Writing – review & editing

Suheyla Cetin-Karayumak: Investigation, Data curation, Writing – review & editing Yang Song: Investigation, Writing – review & editing

Weidong Cai: Investigation, Writing – review & editing

Carl-Fredrik Westin: Investigation, Data curation, Writing – review & editing

James J. Levitt: Investigation, Result Interpretation, Writing – review & editing

Nikos Makris: Investigation, Result Interpretation, Writing – review & editing

Yogesh Rathi: Investigation, Data curation, Writing – review & editing

Lauren J. O’Donnell: Conceptualization, Data curation, Investigation, Project administration, Software, Visualization, Writing – original draft, Writing – review & editing

## Acknowledgment

This work is in part supported by the National Key R&D Program of China (No. 2023YFE0118600) and the National Natural Science Foundation of China (No. 62371107).

## Footnotes

1 The anatomical tract parcellation includes 58 deep white matter fiber tracts, plus 198 short and medium range superficial fiber clusters organized into 16 categories according to the brain lobes they connect.

